# δ-Catenins couple cadherin adhesions to phospholipid-rich membrane domains

**DOI:** 10.64898/2026.08.01.742249

**Authors:** Sergey Ryabichko, Regina B. Troyanovsky, Indrajyoti Indra, Farida V. Korobova, Sergey M. Troyanovsky

## Abstract

δ-Catenins interact with both classical and desmosomal cadherins and play essential, yet incompletely understood, role in adherens junctions (AJs) and desmosomes. According to the prevailing model, δ-catenins are recruited to these junctions exclusively through direct binding to the cadherin juxtamembrane domain (JMD). Here, we show that plakophilin 4 (Pkp4), one of the AJ-associated δ-catenins, is recruited into AJs through two distinct and independent mechanisms. The first is the conventional pathway based on direct interaction with the cadherin JMD. The second is a previously unrecognized mechanism that targets Pkp4 specifically to lateral AJs, submicron-sized, exceptionally stable junctions located along the mid-lateral region of epithelial cell-cell contacts. This targeting occurs independently of the cadherin JMD but requires an interaction with phospholipid-rich plasma membrane domains. We identify the conserved insert between ARM repeats 5 and 6 as the phospholipid-binding module of Pkp4. Because both membrane-binding determinants within this insert, a palmitoylated cysteine residue and a polybasic motif, are highly conserved throughout the δ-catenin family, our findings suggest that recognition of specialized plasma membrane domains is a general property of δ-catenins. We propose that the interplay between cadherin- and phospholipid-dependent targeting mechanisms enables individual δ-catenins to selectively stabilize distinct cadherin-based cell-cell junctions, thereby contributing to the overall architecture of the cell-cell adhesion system.

## Introduction

Adherens junctions (AJs) are critical for overall cell-cell adhesion, cellular architecture, and numerous intercellular signaling pathways (Honig et al., 2020; Mège & Ishiyama, 2017; Lin et al., 2023; Green et al., 2010; Leckband & Rooij, 2014). Their principal adhesive receptors are classical cadherins, which are linked to the actin cytoskeleton through the β-catenin/α-catenin heterodimer to form the cadherin-catenin complex (CCC). A substantial body of evidence indicates that only two core CCC activities, extracellular trans/cis inter-cadherin interactions and intracellular α-catenin oligomerization along actin filaments, are sufficient to generate “minimal” AJs (Troyanovsky, 2023). However, additional CCC-associated activities are required to diversify these “minimal” junctions into highly flexible, regulatable, and cell type-specific adhesive structures. The molecular mechanisms underlying this higher-order regulation remain poorly understood.

One of the principal mediators of this regulation is the δ-catenin (p120-catenin) subfamily of armadillo (ARM) repeat proteins. In mammals, this family comprises four AJ-associated members, p120, plakophilin 4 (Pkp4), δ2-catenin, and ARVCF (Armadillo Repeat Protein Deleted In Velo-Cardio-Facial Syndrome), and three desmosome-associated members, plakophilins 1-3 (Pkp1-3). All seven proteins contain a conserved central ARM-repeat domain flanked by divergent N- and C-terminal regions (Anastasiadis & Reynolds, 2000; Hatzfeld, 2005). While invertebrates generally tolerate loss of their single δ-catenin homolog with minimal phenotypic consequences (Myster et al., 2003; Pettitt et al., 2003), most vertebrate paralogs are largely essential for normal development (McCrea & Park, 2007). Mutations in δ-catenins cause numerous skin, neurological, and cardiac disorders, underscoring their critical functions in both classical cadherin- and desmosomal cadherin-mediated adhesion (Crea & Park, 2007; Bass-Zubek et al., 2009; Hofmann, 2020; Donta et al., 2022).

AJ-associated δ-catenins bind directly to the juxtamembrane domain (JMD) of classical cadherins (Mariner et al., 2000; Anastasiadis & Reynolds, 2000; Ishiyama et al., 2010). This binding suppresses cadherin endocytic motifs located within or adjacent to the JMD, thereby stabilizing CCC at the cell surface (Cadwell et al., 2016; Davis et al., 2003). Whether this function, which is vertebrate-specific, fully accounts for the essential role of δ-catenins in vertebrate AJ organization remains unclear. Current evidence suggests that δ-catenins have broader regulatory functions. For example, the best-characterized family member, p120, is proposed to act as a molecular switch that promotes strong cadherin adhesion in its active state and junctional disassembly in its inactive state (Aono et al., 1999; Anastasiadis & Reynolds, 2001; Oas et al., 2013; Xiao et al., 2007). Likewise, desmosomes fail to assemble in the absence of plakophilins even though desmosomal cadherins are neither destabilized nor degraded (Indra et al., 2021). Proposed mechanisms for δ-catenin function include modulation of the cytoskeleton (Maiden et al., 2016), facilitating CCC oligomerization (Fuchs et al., 2019; Vu et al., 2021), or participation in Rho-family GTPase signaling (Zebda et al., 2013). However, none of these models provides a general concept of how δ-catenins contribute to cell-cell adhesion.

We have recently showed that two AJ-associated δ-catenins, p120 and Pkp4, are essential for distinct AJ subtypes that assemble through fundamentally different mechanisms. Whereas the major cellular δ-catenin, p120, is required for the conventional α-catenin-dependent AJs, referred to here as apical and basal AJs, its minor paralog, Pkp4 is required for a unique α-catenin-independent pathway, forming so-called lateral AJs (Indra et al., 2026). These submicron, dot-like junctions are present along the lateral membrane of nearly all types of epithelial cells and apparently function primarily in cell-cell signaling rather than adhesion (Wu et al. 2014; Indra et al., 2026). A simple possibility is that p120 and Pkp4 stabilize apical/basal and lateral AJs, respectively, by preventing CCC endocytosis specifically at these AJs. A critical unanswered question, however, is what spatial cues direct these proteins to their respective junctions.

Here, we use Pkp4 as a model to define the mechanism by which δ-catenins selectively recognize specific AJ subtypes. We chose Pkp4 because, unlike p120, its complete knockout destabilizes lateral AJs without disrupting overall cell-cell adhesion, greatly simplifying mechanistic analysis. Unexpectedly, we found that Pkp4 associates with lateral AJs even in the absence of direct binding to the cadherin JMD. Instead, this association depends on direct binding of the conserved insert between ARM repeats 5 and 6 to plasma membrane phospholipids, including PI(4,5)P₂. Moreover, this phospholipid-dependent pathway is sufficient to generate trans-interacting Pkp4 clusters at sites of cell-cell contacts even in cadherin-deficient cells. Together, our findings demonstrate that the ARM domain of δ-catenins encodes two parallel targeting mechanisms: one is mediated by direct binding to the cadherin JMD, and the other - by interaction with specialized plasma membrane domains. Although the molecular details of phospholipid-mediated targeting remain to be elucidated, our results suggest that the interplay between cadherin and phospholipid binding provides a general framework by which δ-catenins organize the architecture and specialization of cadherin-based junctions in vertebrate tissues.

## Results

### The cadherin-binding site of Pkp4 is dispensable for Pkp4 association with AJs

Structural and mutagenesis studies (Ishiyama et al., 2010) have shown that the cadherin JMD-binding interface of p120 is formed by a basic groove spanning the αH3 helices of ARM repeats 1-5 (αH3^ARM1/5^). The strong conservation of this groove among AJ-specific δ-catenins suggests that all these proteins use this interface to bind the cadherin JMD. However, the preferences of p120 and Pkp4 for distinct types of AJs (Indra et al., 2026) suggest that these two proteins may also interact with additional partners, which target corresponding CCC complexes to particular AJs. To reveal such additional interactions, we mutated two key cadherin-interacting residues within the αH3^ARM1/5^ groove of GFP-tagged Pkp4, Trp640 and Asn641 (corresponding to Trp477 and Asn478 in p120), to Ala (Fig. 1a). The resulting mutant, GFPpkp4^WN^, and intact GFPpkp4 were expressed in Pkp4-KO A431 cells at comparable levels (Fig. 1b). As expected, co-immunoprecipitation demonstrated that GFPpkp4^WN^, unlike GFPpkp4, exhibited no detectable interactions with E-cadherin (Fig. 1c).

**Figure 1.**
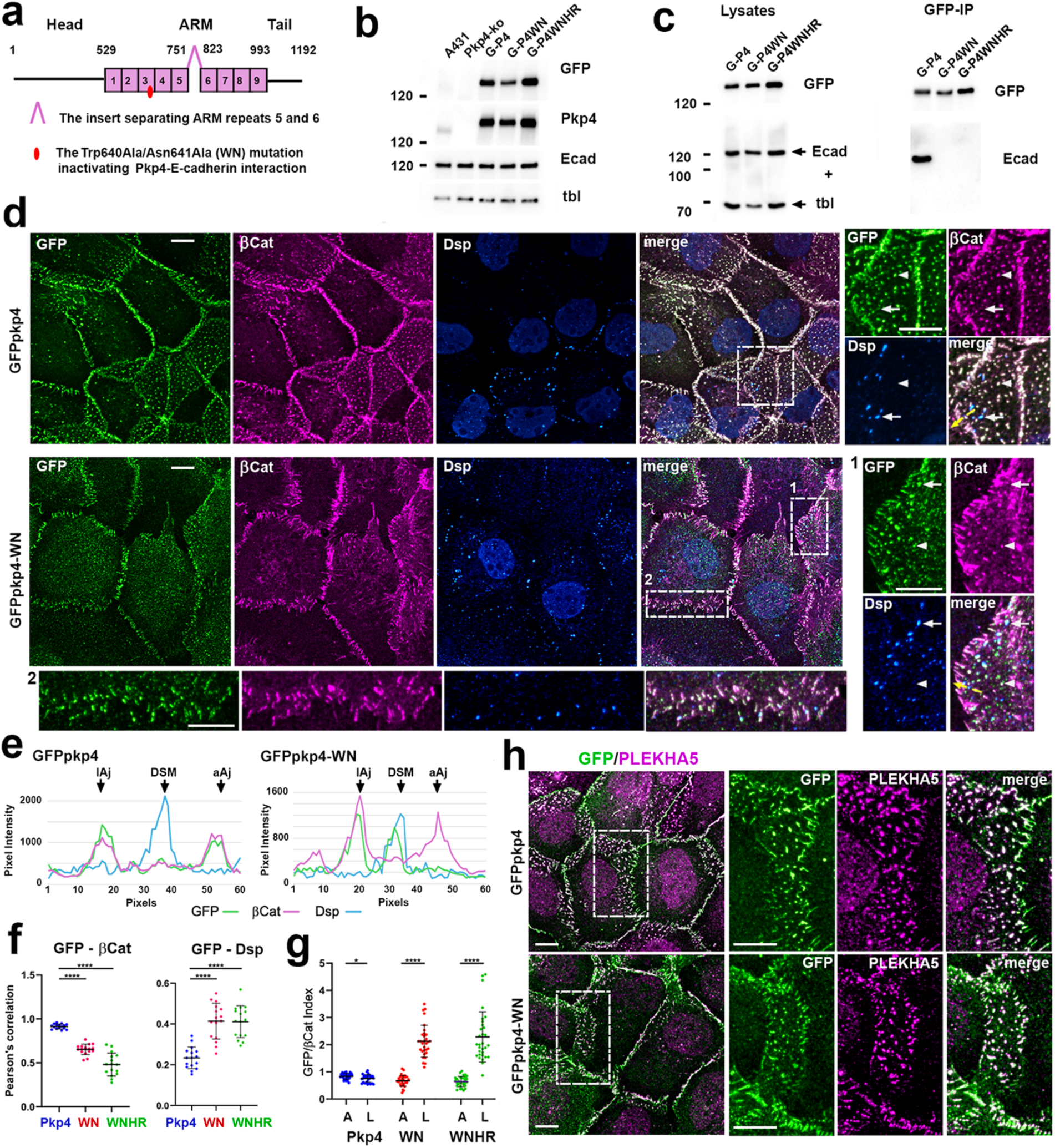
Uncoupling of Pkp4 from classical cadherins does not prevent its association with AJs. (**a**) Schematic representation of Pkp4 showing the N-terminal head, ARM repeat, and C-terminal tail domains. Individual ARM repeats are numbered. The ARM domain is interrupted by a 72 amino acid insert between repeats 5 and 6 (INT5/6). Amino acid boundaries of each domain are indicated. (**b**) Immunoblot analysis of wild type A431 cells (A431), the A431-Pkp4-KO subline (Pkp4-ko), and its derivatives expressing GFPpkp4 (G-P4), GFPpkp4^WN^ (G-P4WN), or GFPpkp4^WNHR^ (G-P4WNHR). Blots were probed for GFP, endogenous Pkp4 (Pkp4), E-cadherin (Ecad) and β-tubulin staining (tbl; loading control). Molecular mass markers (kDa) are on the left. (**c**) Representative co-immunoprecipitation experiment. Total cell lysates (Lysates) before anti-GFP immunoprecipitation and the resulting precipitates (GFP-IP) were immunoblotted for GFP (GFP) and E-cadherin/tubulin (Ecad+tbl). Note, both Pkp4 mutants failed to co-immunoprecipitate E-cadherin. (**d**) Maximum-intensity projections of confocal z-stacks from cells expressing GFPpkp4 or GFPpkp4^WN^ stained for GFP, β-catenin (βCat), and desmoplakin (Dsp). Boxed regions are enlarged to the right or below. Scale bars, 10 μm. Note GFPpkp4 was similarly enriched at all β-catenin-positive AJs. In contrast, GFPpkp4^WN^ preferentially accumulated at lateral AJs (arrowheads) but not apical AJs. Arrows indicate desmosomes, which lacked GFPpkp4 but contained GFPpkp4^WN^. (**e**) Fluorescence intensity profiles measured along the dashed lines shown in (**d**). Note that GFP and β-Catenin fluorescence coincide at both lateral and apical AJs (lAj and aAj, respectively) in GFPpkp4 cells, but are present only in lAJ in the mutant-expressing cells. GFPpkp4^WN^ is also detected at desmosomes. (**f**) PCC for GFP/β-catenin and GFP/desmoplakin fluorescence within cell-cell contacts of cells expressing GFPpkp4, GFPpkp4^WN^, or GFPpkp4^WNHR^. (**g**) GFP/β-catenin fluorescence ratio (GFP/βCat index) in apical (A) and lateral (L) AJs. Note, in GFPpkp4-expressing cells, this ratio was approximately 1 for both AJ subtypes. In contrast, the ratio increased significantly in lateral AJs of cells expressing GFPpkp4^WN^ or GFPpkp4^WNHR^, reflecting preferential enrichment of the cadherin-uncoupled mutants at lateral AJs. Statistical significance in (**f** and **g**) was calculated using two-tailed Student’s t tests: *P < 0.05; ****P < 0.0001. The means ± SD are indicated by bars. (**h**) Maximum-intensity projections of confocal z-stacks from cells shown in (**d**) stained for GFP and PLEKHA5. Merged images are shown on the left. Boxed regions are enlarged on the right and presented in separate fluorescence channels (scale bars, 10 μm).

To characterize the subcellular distribution of the GFP-tagged proteins, the cells were stained for GFP, in combination with obligate AJ and desmosomal markers, β-catenin, and desmoplakin (Dsp), correspondingly. The later marker was included since Pkp4 or its mutants had been detected in desmosomes (Calkins et al., 2003; Hatzfeld et al., 2003). Supporting our idea, this staining showed that despite the loss of cadherin binding, GFPpkp4^WN^ still formed clusters at cell-cell contacts. However, its co-localization with β-catenin was reduced, as reflected by a decrease in GFP/β-catenin Pearson’s correlation coefficient (PCC) from ∼0.9 to ∼0.7 (Fig. 1e). At the same time, GFPpkp4^WN^ displayed clear association with desmosomes, whereas intact GFPpkp4 was restricted almost exclusively to AJs (see PCC and line scans in Figs. 1f and e). The most striking feature of GFPpkp4^WN^ was its strong enrichment in lateral AJs, some of which contained only barely detectable β-catenin signal (Fig. 1d and line scan, Fig 1e). Staining for the lateral AJ marker PLEKHA5 confirmed the identity of these structures as lateral AJs (Fig. 1h). To quantify the GFPpkp4^WN^ preference to lateral versus apical AJs, we determined their GFP/β-catenin fluorescence ratio (GFP/βCat index). Whereas this index was nearly identical for both apical and lateral AJs of GFPpkp4-expressing cells (∼0.9), it diverged markedly in GFPpkp4^WN^-expressing cells, increasing to ∼2.1 for lateral AJs and, in contrast, reducing to ∼0.7 for apical AJs (Fig. 1g).

To exclude the possibility that residual, undetectable by co-IP assay cadherin binding accounted for AJ localization of Pkp4^WN^, we introduced into this mutant two additional charge-reversing mutations (H554E, R595E) in its αH3^ARM1/5^ groove. AlphaFold3 structural modeling predicted that the resulting Pkp4^WNHR^ mutant was unable to dock the cadherin JMD into its αH3^ARM1/5^ groove (Fig. S1a). Nevertheless, GFPpkp4^WNHR^ was indistinguishable from GFPpkp4^WN^ by all tested characteristics, including preferential localization to lateral AJs, incorporation into PLEKHA5-positive structures, and increased association with desmosomes (Figs. 1f,g and S1b–e). Together, these results demonstrate that in addition to the direct cadherin binding, which recruits the overexpressed GFPpkp4 into all types of AJs, Pkp4 contains another, cadherin-independent determinant, which specifically targets GFPpkp4^WN^ mutant into lateral AJs and desmosomes.

### Pkp4 clusters are cadherin-independent and interact across the cell-cell contacts

To determine whether any AJ-associated proteins mediate Pkp4 recruitment into AJs, we expressed GFPpkp4^WN^ in cadherin-deficient A431-Ec/Pc-KO cells (Troyanovsky et al., 2021). Because these cells lack classical cadherins and do not form AJs, we expected GFPpkp4^WN^ to accumulate diffusely in the cytoplasm. Surprisingly, the mutant formed numerous clusters throughout the cells, a subset of which localized to cell-cell contacts. Co-staining for β-catenin, which associates with all type I and type II cadherins, confirmed that these contact-associated clusters did not contain detectable amounts of any cadherin species, which might remain in Ec/Pc-KO cells as minor components (Fig. 2a). Likewise, staining for the desmosomal cadherin, desmoglein 2 (Dsg2), demonstrated that most clusters were not desmosomes (Fig. 2a). Finally, only a small fraction of the clusters recruited the lateral AJ marker PLEKHA5 (Fig. 2a). Together, these observations indicate that neither classical nor desmosomal cadherins, nor PLEKHA5, are required for Pkp4 cluster formation or for their localization to cell-cell contacts. Next, we asked whether these cadherin-independent Pkp4 clusters could interact in trans. To this end, we co-cultured two A431-Ec/Pc-KO clones, one expressing GFPpkp4^WN^ and the other expressing the same mutant fused to a red fluorescent protein, mCherry (mCHpkp4^WN^). The co-cultures showed numerous green and red clusters aligned precisely across heterochromatic cell-cell contacts (Figs. 2b,c). These aligned clusters were negative for desmoplakin, excluding the possibility that they represented desmosomes. Thus, despite the absence of major classical cadherins and obliteration of the cadherin-binding interface that makes unlikely the interaction even with minor cadherin species, Pkp4 clusters retained the ability to recognize one another across apposing plasma membranes and to form trans-interacting structures.

**Figure 2.**
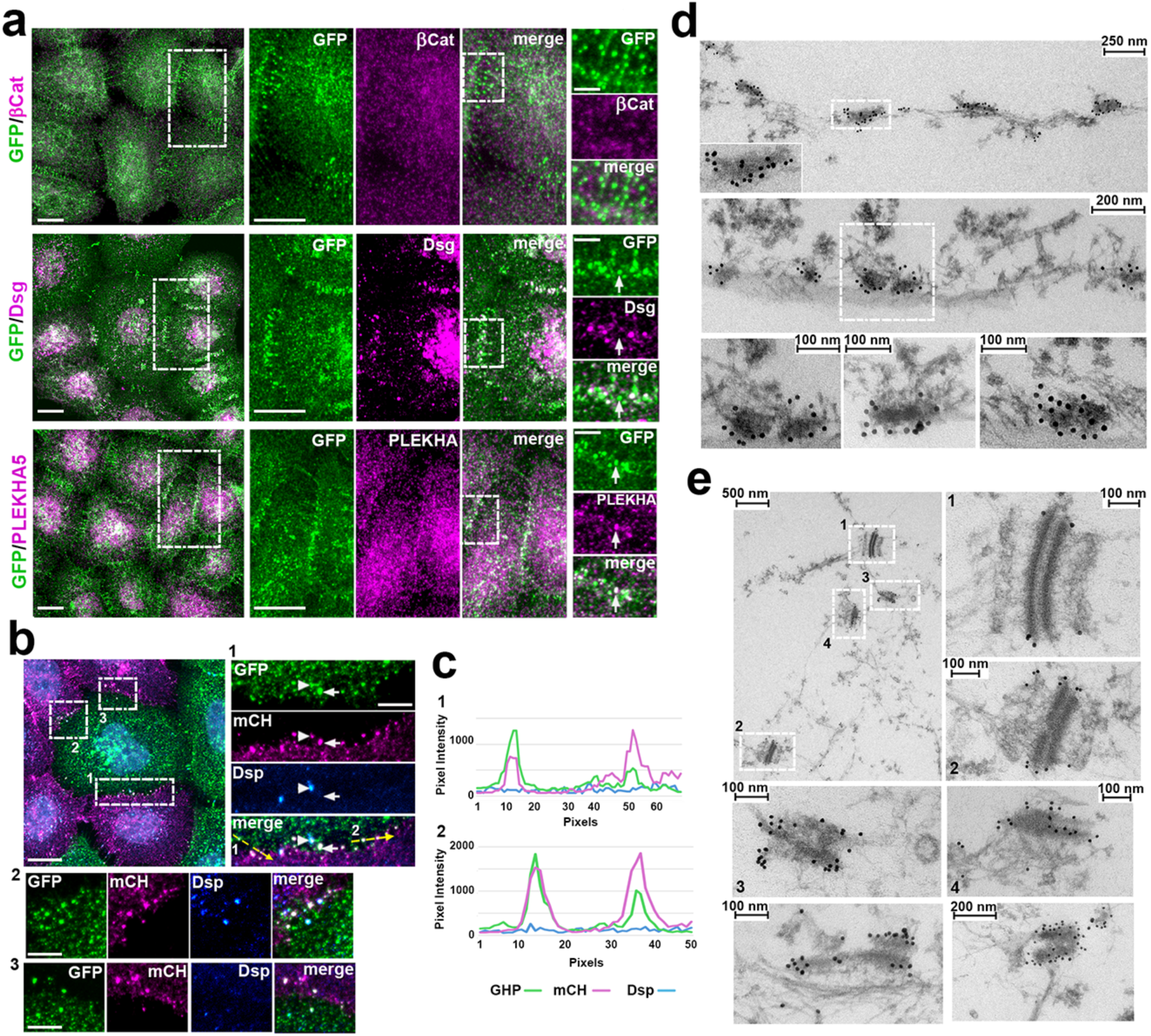
Cadherin-deficient Pkp4 clusters interact in trans. (**a**) Maximum-intensity projections of confocal z-stacks from GFPpkp4^WN^-expressing A431-Ec/Pc-KO cells stained for GFP together with β-catenin (GFP/βCat), desmoglein 2 (GFP/Dsg), or PLEKHA5 (GFP/PLEKHA5). Low-magnification merged images are shown on the left. Boxed regions are enlarged in the center as separate fluorescence channels (scale bars, 10 μm), and selected regions are further magnified on the right (scale bars, 2.5 μm). GFPpkp4^WN^ clusters were uniformly negative for β-catenin, confirming the absence of classical cadherins. Only a small subset of clusters was positive for Dsg or PLEKHA5. (**b**) Co-culture of A431-Ec/Pc-KO cells expressing GFPpkp4^WN^ or mCHpkp4^WN^. Cells were stained for GFP (green) and mCherry (magenta) and Dsp (blue). The merged low-magnification image is shown on the left (scale bar, 10 μm). Three representative heterochromatic cell-cell contacts, outlined by numbered dashed boxes (numbered), are enlarged on the right and below (scale bar, 3 μm) and shown as individual fluorescence channels. Dsp-negative but GFP/mCherry-positive clusters (one indicated by an arrow) are precisely aligned across the cell-cell contacts, indicating that cadherin-independent Pkp4 clusters interact in trans. One of the desmosomes is indicated by an arrowhead. (**c**) Fluorescence intensity profiles measured along the numbered dashed yellow lines shown in (**b**). Note that GFP and mCH fluorescence profiles closely coincide despite significant variations in red and green fluorescence intensities. No Dsp signal is detected. (**d**) TEM of plasma membrane-associated nanogold-labeled GFPpkp4^WN^ clusters located outside of cell-cell contact regions. Boxed regions are zoomed in the inset or at the bottom left. Two additional high-magnification examples from other cells are shown on the right. (**e**) TEM of cell-cell contact-associated structures. A representative cell-cell contact region is shown at low magnification (upper left). Four junctional structures are numbered: two desmosomes (1 and 2) and two GFP-labeled GFPpkp4^WN^-enriched junctions (3 and 4). High-magnification views of the numbered structures are shown to the right and below. Two additional examples of GFPpkp4^WN^-enriched junctions from other cells are shown in the bottom row.

To further characterize both junctional and non-junctional GFPpkp4^WN^ clusters, we performed their anti-GFP immunogold labeling followed by transmission electron microscopy (TEM). To preserve the cytoskeleton while ensuring antibody accessibility, cells were extracted with 1% Triton X-100 in cytoskeleton-preserving buffer before fixation, as described previously (Indra et al., 2026). Most non-junctional GFPpkp4^WN^ clusters appeared as electron-dense structures associated with both the plasma membrane and the cytoskeleton (Fig. 2d). Cell-cell contacts of these cells contained typical desmosomes (Fig. 2e), readily identified by their associated intermediate filament bundles and symmetric electron-dense outer plaques separated by a relatively uniform intercellular space (20–25 nm). In most desmosomes, gold particles were detected only at the plaque periphery. In addition, cell-cell contacts contained heavily labeled junctional structures that likely corresponded to the cadherin-independent Pkp4^WN^ clusters. These structures also exhibited electron-dense outer plaques, which, however, differed markedly from desmosomes: the plaques showed only weak association with the cytoskeleton, they were highly irregular in thickness (5–50 nm) and most importantly, nearly always asymmetric between the two apposing membranes. This was consistent with the high variability of the red and green fluorescence intensities of the individual junctions in the co-cultures described above (see line scans in Fig. 2c). Together, our observations demonstrate that cadherin-uncoupled Pkp4, even in cells lacking major classical cadherins, forms membrane-associated electron-dense assemblies, a subset of which establishes distinctive trans-interacting junctional structures. While one cannot exclude a possibility that these junctional structures are initiated by remaining minor cadherin species, these observations imply that a bulk of Pkp4 is delivered to the plasma membrane by non-cadherin-based mechanisms.

### The 5/6 insert within the ARM domain mediates interactions with a specific set of phospholipids

We recently showed that the ARM domain of desmosomal δ-catenin Pkp3 (ARM^PKP3^) interacts with both actin filaments and phospholipids (Gupta et al., 2024). Either of these two interactions could potentially promote the cadherin-independent recruitment of Pkp4 to cell-cell contacts, which are known to be enriched in specific phospholipid domains (Xiong et al., 2012; Kanemaru et al. 2022) and associated with distinct cortical actin networks (DeMali & Burridge., 2003; Troyanovsky et al., 2025). To determine whether the ARM domain of Pkp4 (ARM^PKP4^) also interacts with F-actin and/or phospholipids, we probed these interactions by in vitro binding assays using recombinant GST-tagged ARM^PKP4^ (GST-ARM^PKP4^). Actin co-sedimentation assay detected only weak interactions of ARM^PKP4^ with F-actin (Fig. S2). In contrast, lipid-overlay assay with Echelon Mega Lipid Strips containing 23 major membrane lipids, revealed robust binding of ARM^PKP4^ to several phospholipids, including PI(4,5)P2 and PI(3,4)P2, two phosphoinositides previously detected in cell-cell contacts (Fig. 3a).

**Figure 3.**
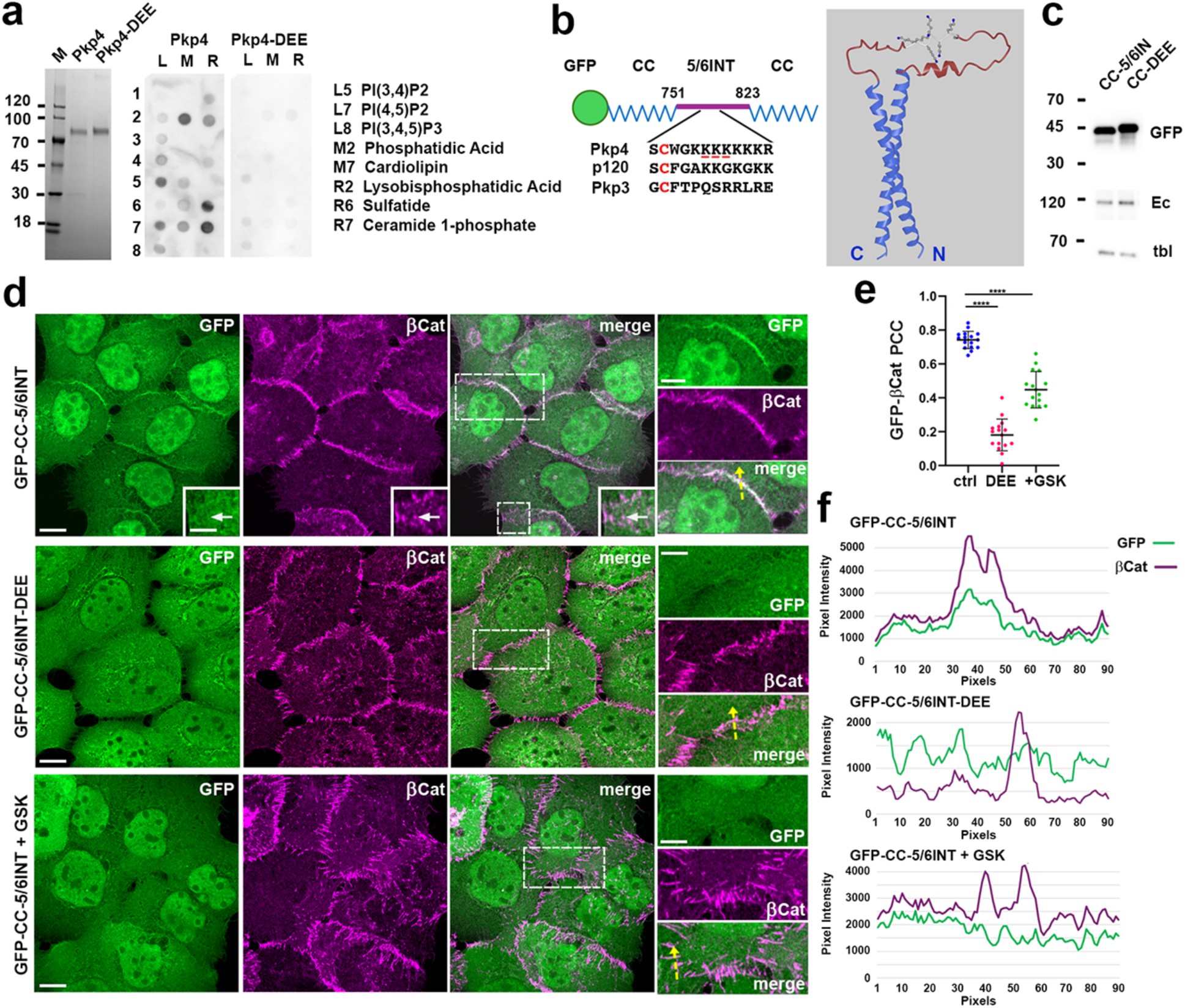
The 5/6 insert of Pkp4 interacts with phospholipid-rich plasma membrane domains. (**a**) Recombinant GST-tagged ARM domain of Pkp4 (Pkp4) or its K788D/K789E/K790E mutant (Pkp4-DEE) were analyzed by SDS-PAGE (left) and for phospholipid binding using Echelon Mega Lipid Strips (center). Phospholipids showing the strongest binding signals are listed on the right. (**b**) Schematic representation (left) of the GFP-CC-5/6INT^PKP4^ biosensor, consisting of GFP (green) fused to a two-stranded antiparallel coiled-coil module (CC, blue) derived from seryl-tRNA synthetase (Amer et al., 2023) with the Pkp4 5/6INT inserted into the loop connecting the two helices. The central region of the 5/6 insert containing the conserved cysteine residue (red) and the polybasic motifs of Pkp4, p120, and Pkp3 are shown below. The AlphaFold model of the biosensor (without GFP) is shown on the right, with residues of the polybasic motif displayed as ball-and-stick representations. (**c**) Immunoblot analysis of A431-Pkp4-KO cells expressing GFP-CC-5/6INT^PKP4^ (CC-5/6INT) or its DEE mutant (CC-DEE). Blots were probed for GFP, E-cadherin (Ec), and β-tubulin (Tbl; loading control). (**d**) Maximum-intensity projections of all x-y optical slices of cells expressing GFP-CC-5/6INT^PKP4^ (GFP-CC-5/6INT), its DEE mutant (GFP-CC-5/5INT-DEE), or GFP-CC-5/6INT-treated for 30 min with the PI4KIIIα inhibitor GSK-A1 (GFP-CC-5/6INT + GSK). Cells were stained for GFP and β-catenin (βCat). Scale bars, 10 μm. Boxed regions are enlarged on the right or in the insets. Scale bars, 5 μm. GFP-CC-5/6INT was enriched along cell-cell contacts but did not preferentially accumulate at lateral AJs (one lateral AJ is indicated by an arrow). In contrast, the DEE mutation or GSK-A1 treatment abolished this cell-cell contact localization and redistributed the biosensor to the cytoplasm. (**e**) Pearson’s correlation coefficients for GFP and β-catenin fluorescence within cell-cell contacts of cells expressing the biosensor (Ctrl), its DEE mutant (DEE), or treated with GSK-A1 (GSK). Statistical significance was calculated using two-tailed Student’s t tests: ****P < 0.0001. The means ± SD are indicated by bars. (**f**) Fluorescence intensity profiles measured along the dashed yellow lines shown in (d). GFP and β-catenin fluorescence largely coincide at cell-cell contacts in cells expressing GFP-CC-5/6INT, whereas little correlation is observed after introduction of the DEE mutation or treatment with GSK-A1.

To identify the site responsible for phospholipid binding in our assay, we examined ARM^PKP4^ for candidate membrane-interacting sites. A prominent feature of all δ-catenins is a relatively conserved insert separating ARM repeats 5 and 6. In Pkp4, this 72-amino-acid segment (5/6INT^PKP4^) contains two potential lipid-binding determinants: a conserved cysteine residue (Cys785), which was shown to undergo palmitoylation in desmosomal δ-catenins (Roberts et al., 2014), and a polybasic motif (KKKKKKKR; residues 788–795) potentially capable of electrostatic interactions with acidic phospholipids (Fig. 3b). To test the role of this site in lipid binding, we generated an GST-ARM^PKP4-DEE^ mutant carrying three charge-reversing substitutions, K788D, K789E, and K790E (DEE mutation, underlined in Fig. 3b) within the polybasic motif. This mutant showed markedly reduced phospholipid binding in the overlay assay (Fig. 3a), indicating that the polybasic motif contributes to phospholipid recognition.

We next asked whether this polybasic motif is sufficient to target Pkp4 to cell-cell contacts. To this end, we generated a biosensor (GFP-CC-5/6INT^PKP4^) by inserting the entire 5/6INT^PKP4^ segment into a GFP-tagged coiled-coil module derived from seryl-tRNA synthetase (Fig. 3b). When expressed in A431 Pkp4-KO cells, the biosensor accumulated efficiently at cell-cell contacts and frequently co-localized with β-catenin (Fig. 3c,d), yielding a high PCC (∼ 0.8) and overlapping fluorescence profiles across cell-cell contacts (Fig. 3e,f). Despite this cell-cell contact targeting, the biosensor did not display notable enrichment in lateral AJs (see a representative lateral AJ marked by the arrow in Fig. 3d). Importantly, the DEE mutations of the polybasic motif completely abolished cell-contact localization of the biosensor, resulting in diffuse cytoplasmic distribution and a significant reduction in co-localization with AJs (Fig. 3c–f).

Since PI(4,5)P₂ and PI(3,4)P₂ were among the strongest phospholipid ligands of GST-ARM^PKP4^, we next examined the role of phosphoinositides in biosensor targeting. Cells expressing GFP-CC-5/6INT^PKP4^ were treated with GSK-A1, a potent and selective inhibitor of PI4KIIIα, the kinase directly responsible for plasma membrane production of PI(4)P and, indirectly, for entire phospholipid composition of plasma membrane (Bojjireddy et al., 2014; Burke et al., 2023; Huang et al., 2026). Remarkably, a 30-min treatment completely displaced the biosensor from the plasma membrane, as evidenced by a pronounced decrease in β-catenin-GFP PCC and an absence of the overlaps with E-cadherin in the line scans (Fig. 3e,f). Taken together, these findings identify the 5/6INT^PKP4^ as a phospholipid-binding module of Pkp4.

### Interactions with plasma membrane lipids drive cadherin-independent Pkp4 clustering at cell-cell contacts

To determine whether phospholipid binding is required for targeting Pkp4 to cell-cell contacts, we introduced lipid-uncoupling mutations into GFPpkp4^WN^. In one mutant, GFPpkp4^WN-C785A^, substitution of the conserved Cys785 with Ala was expected to prevent potential Pkp4 palmitoylation. In the second mutant, GFPpkp4^WN-DEE^, the DEE mutations disrupted the phospholipid-binding polybasic motif described above. These constructs were expressed in A431 Pkp4-KO cells at levels comparable to those of GFPpkp4^WN^ (Fig. 4a). Triple staining of these cells for GFP, AJs and desmosomes showed that both mutations produced nearly identical phenotypes (Fig. 4b). They almost completely abolished the association of GFPpkp4^WN^ clusters with any type of AJs marked by β-catenin (Fig. 4b, arrowheads). Few remaining contact-associated mutant clusters corresponded almost exclusively to desmosomes (Fig. 4b, arrows). Consistent with these observations, the PCC between Pkp4 and β-catenin was reduced from approximately 0.5 in the GFPpkp4^WN^ cells to ∼0.2 in GFPpkp4^WN-DEE^ cells (Fig. 4c). An even stronger effect was revealed by quantifying the AJ enrichment coefficient of the mutants, defined as the ratio of junctional to extra-junctional GFP fluorescence. In control cells, β-catenin-marked AJs contained approximately six-fold more GFPpkp4^WN^ fluorescence than the surrounding membrane. This value decreased to approximately two-fold in cells expressing either GFPpkp4^WN-DEE^ or GFPpkp4^WN-C785A^ (Fig. 4d). No changes in this parameter were detected for β-catenin. Thus, interactions with phospholipids are required for efficient clustering of Pkp4^WN^ within AJs.

**Figure 4.**
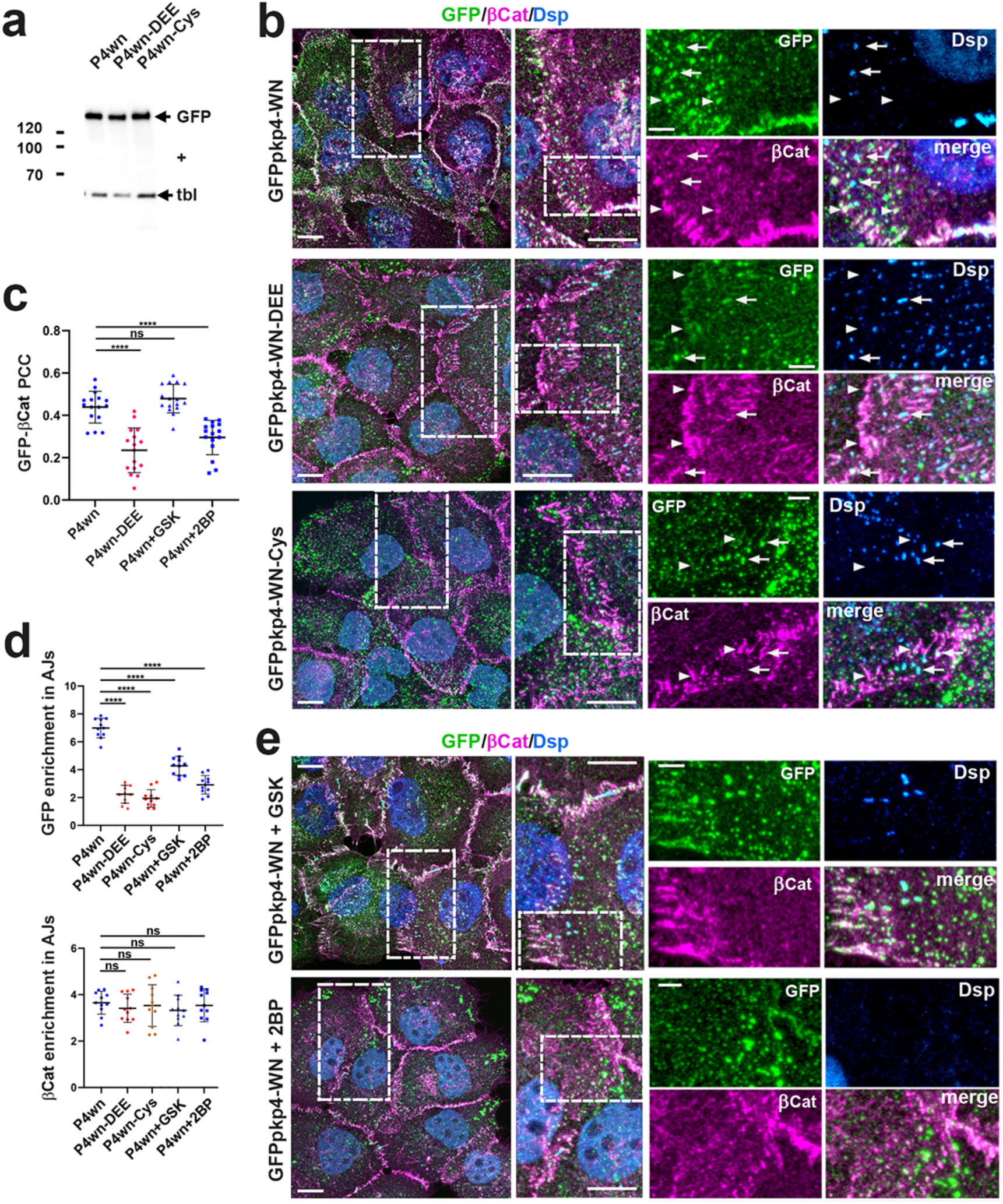
Interactions with phospholipids are required for cadherin-independent clustering of Pkp4 at cell-cell contacts. (**a**) Comparative immunoblot analysis of A431-Pkp4-KO cells expressing the cadherin-uncoupled GFPpkp4^WN^ (P4wn), and its derivatives, GFPpkp4^WN-DEE^ and GFPpkp4^WN-C785A^ (P4wn-DEE and P4wn-Cys, respectively) carrying additional mutations either in the phospholipid-binding site or in the conserved palmitoylated cysteine. Blots were probed for GFP and β-tubulin (tbl; loading control). Molecular mass markers (kDa) are on the left. (**b**) Maximum-intensity projections of confocal z-stacks from cells expressing GFPpkp4^WN^ (GFPpkp4-WN), GFPpkp4^WN-DEE^ (GFPpkp4-WN-DEE), and GFPpkp4^WN-C785A^ (pkp4-WN-Cys). Cells were stained for GFP (green), β-catenin (βCat, magenta), and Dsp (blue). Representative cell-cell contact regions (dashed boxes) are zoomed on the right. Scale bars, 10 μm. Boxed areas of the enlarged images are shown at higher magnification as individual fluorescence channels. Scale bars, 2.5 μm. Disruption of phospholipid binding by either the DEE or C785A mutation markedly reduced localization of GFP-Pkp4WN to AJs (arrowheads), whereas localization to desmosomes (marked by arrows) was largely preserved. (**c**) Pearson’s correlation coefficients for GFP and β-catenin fluorescence within cell-cell contacts of cells expressing GFPpkp4^WN^ and its DEE mutant (P4wn and P4wn-DEE, correspondingly), as well as GFPpkp4^WN^-expressing cells treated with GSK-A1 (+GSK) or the palmitoylation inhibitor, 2-bromopalmitate (+2BP). (**d**) Enrichment of GFP-tagged Pkp4 mutants at AJs, expressed as the ratio of junctional to extrajunctional GFP fluorescence. As a control, the corresponding β-catenin enrichment is shown. Mutant designations and treatment conditions are the same as in (**a**) and (**c**). Whereas β-catenin enrichment was unaffected, Pkp4 enrichment at AJs was markedly reduced by disruption of phospholipid interactions, either by mutation or by pharmacological inhibition. Statistical significance in (**c** and **d**) was calculated using two-tailed Student’s t tests: ns, nonsignificant; ****P < 0.0001. The means ± SD are indicated by bars. (**e**) Maximum-intensity projections of confocal z-stacks from cells expressing GFPpkp4^WN^ treated with GSK-A1 or 2-bromopalmitate (GFPpkp4-WN+GSK or GFPpkp4-WN+2BP, respectively). Cells were stained and displayed as in (**b**). Note the pronounced reduction in GFPpkp4^WN^ localization at AJs following 2-bromopalmitate treatment.

We next asked whether acute pharmacological disruption of Pkp4-lipid interactions would similarly affect AJ-associated GFPpkp4^WN^ clusters. In contrast to the biosensor experiments, a 30-min treatment with the PI4KIIIα inhibitor GSK-A1 produced no obvious visual changes in GFPpkp4^WN^ localization (Fig. 4e,c), although it caused a modest but statistically significant decrease in the AJ enrichment coefficient (Fig. 4d). We reasoned that depletion of phosphoinositides alone may be insufficient to fully detach GFPpkp4^WN^ from the membrane because the protein remains membrane-associated through palmitoylation. To test this possibility, GFPpkp4^WN^-expressing cells were treated for 4 h with 2-bromopalmitate (2BP), an irreversible inhibitor of protein acyltransferases that blocks protein palmitoylation (Davda et al., 2013). In contrast to GSK-A1, 2BP treatment almost completely eliminated AJ-associated GFPpkp4^WN^ clusters (Fig. 4e) and dramatically reduced both the GFP-β-catenin PCC value (Fig. 4c) and the AJ enrichment coefficient of the mutant (Fig. 4d). These findings provide additional evidence that the association of GFPpkp4^WN^ clusters with AJs does not depend on residual interactions with classical cadherins. Rather, it is driven by interactions with specialized AJ-associated plasma membrane domains.

### The ARM domain is sufficient to target Pkp4 into lateral AJs

Previous studies have shown that the N- and C-terminal domains of plakophilins, including Pkp4, interact with multiple desmosomal proteins, such as desmosomal cadherins, Dsp, and plakoglobin (Hatzfeld et al., 2003). These interactions, particularly those involving plakoglobin, could potentially contribute to the localization of the cadherin-uncoupled GFPpkp4^WN^ mutant to cell-cell contact adhesions including AJs and desmosomes. To test this mechanism, we generated a Pkp4 mutant lacking both the N- and C-terminal domains, GFPpkp4ΔN/C^WN^ (Fig. 5a), and expressed it in Pkp4-KO A431 cells. As expected, co-immunoprecipitation experiments showed that GFPpkp4ΔN/C^WN^ did not interact with E-cadherin (Fig. 5b). Furthermore, in contrast to full-length GFPpkp4^WN^, GFPpkp4ΔN/C^WN^ failed to form prominent AJ-associated clusters suggesting that the deleted N- and C-terminal domains are required for cluster formation. Nevertheless, consistent with the phospholipid-binding activity of the 5/6INT^PKP4^, the mutant was efficiently targeted to cell-cell contacts, where it displayed a distribution resembling the localization pattern of the 5/6INT^PKP4^ biosensor, e.g., partially co-localized with β-catenin (Fig. 5c). However, in the striking difference from the biosensor distribution, the GFPpkp4ΔN/C^WN^ mutant retained a strong preference for lateral AJs (see arrows in Fig. 5c). Consistently, line scans through cell-cell contacts showed that its overlap with lateral AJs was considerably more pronounced than its association with apical or basal AJs (Fig. 5d).

**Figure 5.**
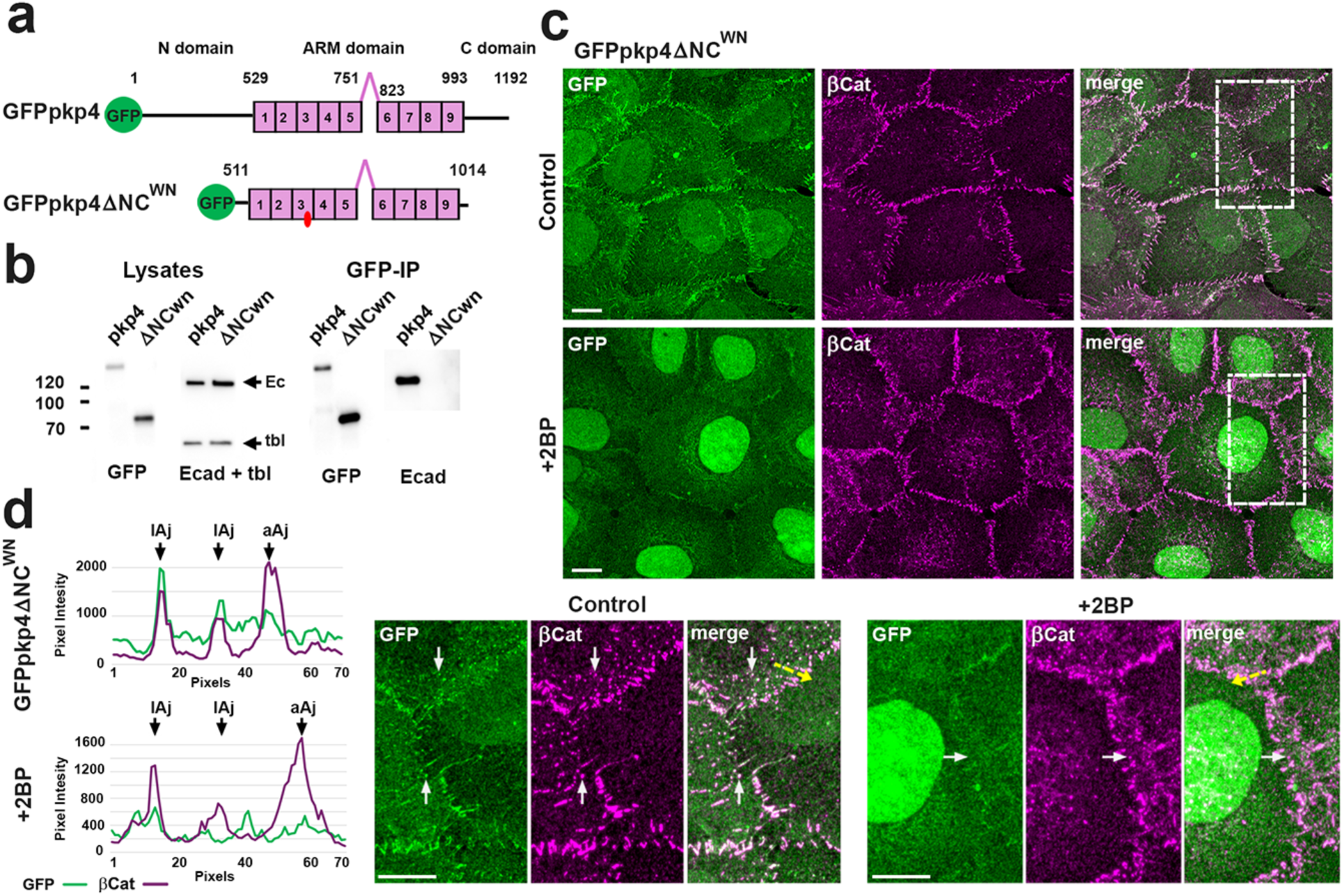
Cadherin-uncoupled ARM domain of Pkp4 retains targeting to lateral AJ. (**a**) Schematic representation of the GFPpkp4 ΔNC^WN^ mutant compared with the full-length Pkp4. All abbreviations are as in Fig. 1a. The mutant lacks both the N-terminal (head) and C-terminal (tail) domains of Pkp4. (**b**) Representative co-immunoprecipitation experiment with GFPpkp4 (Pkp4) and GFPpkp4ΔNC^WN^ (ΔNCwn). Total cell lysates before anti-GFP immunoprecipitation (Lysates) and the resulting immunoprecipitates (GFP-IP) were immunoblotted for GFP and E-cadherin/tubulin (Ecad+tbl). Note, that the GFPpkp4ΔNC^WN^ fails to co-immunoprecipitate E-cadherin. (**c**) Maximum-intensity projections of confocal z-stacks from cells expressing GFPpkp4ΔNC^WN^ under control conditions (Control) or after treatment with 2-bromopalmitate (+2BP). Cells were stained for GFP (green) and β-catenin (βCat, magenta). Representative cell-cell contact regions (dashed boxes) are enlarged below. Several lateral AJs are indicated by arrows. Note the pronounced reduction in GFPpkp4ΔNC^WN^ localization at lateral AJs following 2-bromopalmitate treatment. Scale bars, 10 μm. (**d**) Fluorescence intensity profiles measured along the dashed yellow lines shown in (c). Under control conditions, GFP and β-catenin fluorescence peaks coincide at lateral AJs, whereas this correlation is markedly reduced after 2BP treatment.

Treatment with 2BP strongly diminished plasma membrane localization of GFPpkp4ΔN/C^WN^ (Figs. 5d,e). Unexpectedly, instead of accumulating in cytoplasmic aggregates, as observed for Pkp4^WN^, this treatment redirected GFPpkp4ΔN/C^WN^ to the nucleus (Fig. 5e). Taken together, these results suggest a structure-function framework for Pkp4. Its N- and C-terminal domains are required for efficient cluster formation, whereas the ARM domain contains at least three distinct activities: binding to cadherins, binding to phospholipids, and selective targeting to lateral AJs. The third activity, however, depends on phospholipid binding. In addition, the ARM domain appears to harbor a nuclear-targeting activity that becomes apparent when both N/C domains and membrane association are disrupted.

### Membrane-uncoupled Pkp4 mutants exhibit only a mild phenotype in steady-state cultures

We next examined whether phospholipid binding influences the overall organization of AJs and contributes to the incorporation of Pkp4-containing CCC (Pkp4-CCC) into AJs. To this end, we inserted the membrane-uncoupling mutation DEE into GFP-tagged Pkp4. To prevent potential compensation by other AJ-specific δ-catenins, both, the resulting mutant, GFPpkp4^DEE^, and the control, GFPpkp4, were expressed in δCat-KO A431 cells, which lack p120, Pkp4, and ARVCF (Indra et al., 2026). Clones expressing comparable levels of GFPpkp4 and GFPpkp4^DEE^ were selected (Fig. 6a) and triple-stained for GFP, β-catenin, and Dsp (Fig. 6b). Surprisingly, the DEE mutation produced no obvious defects in AJ assembly or the overall AJ organization: based on β-catenin staining, GFPpkp4^DEE^-expressing cells assembled an AJ network that was indistinguishable from that of control GFPpkp4-expressing cells. However, localization of the mutant exhibited two striking abnormalities. First, unlike wild-type GFPpkp4, which was excluded from desmosomes, GFPpkp4^DEE^ was readily incorporated into these adhesions (Fig. 6b, arrows, and 6c). Second, mutant-expressing cells accumulated numerous GFP-positive, β-catenin-negative intracellular aggregates, particularly common along free cell edges (Fig. 6b, arrowheads). Such structures were not observed in cells expressing wild-type GFPpkp4. Because both desmosomes and the intracellular aggregates lacked β-catenin, these changes resulted in a modest but statistically significant decrease in the PCC between GFP and β-catenin (Fig. 6d). A more pronounced difference was revealed by quantifying of the pool of GFP-positive pixels lacking β-catenin fluorescence, which showed a marked increase in the mutant-expressing cells (Fig. 6d). Taken together, these findings indicate that direct binding to cadherin is sufficient to recruit Pkp4 into AJs. However, Pkp4-phospholipid binding functions to spatially restrict Pkp4 clustering to AJs eliminating its potential to form ectopic clusters associated with desmosomes or in cytosol.

**Figure 6.**
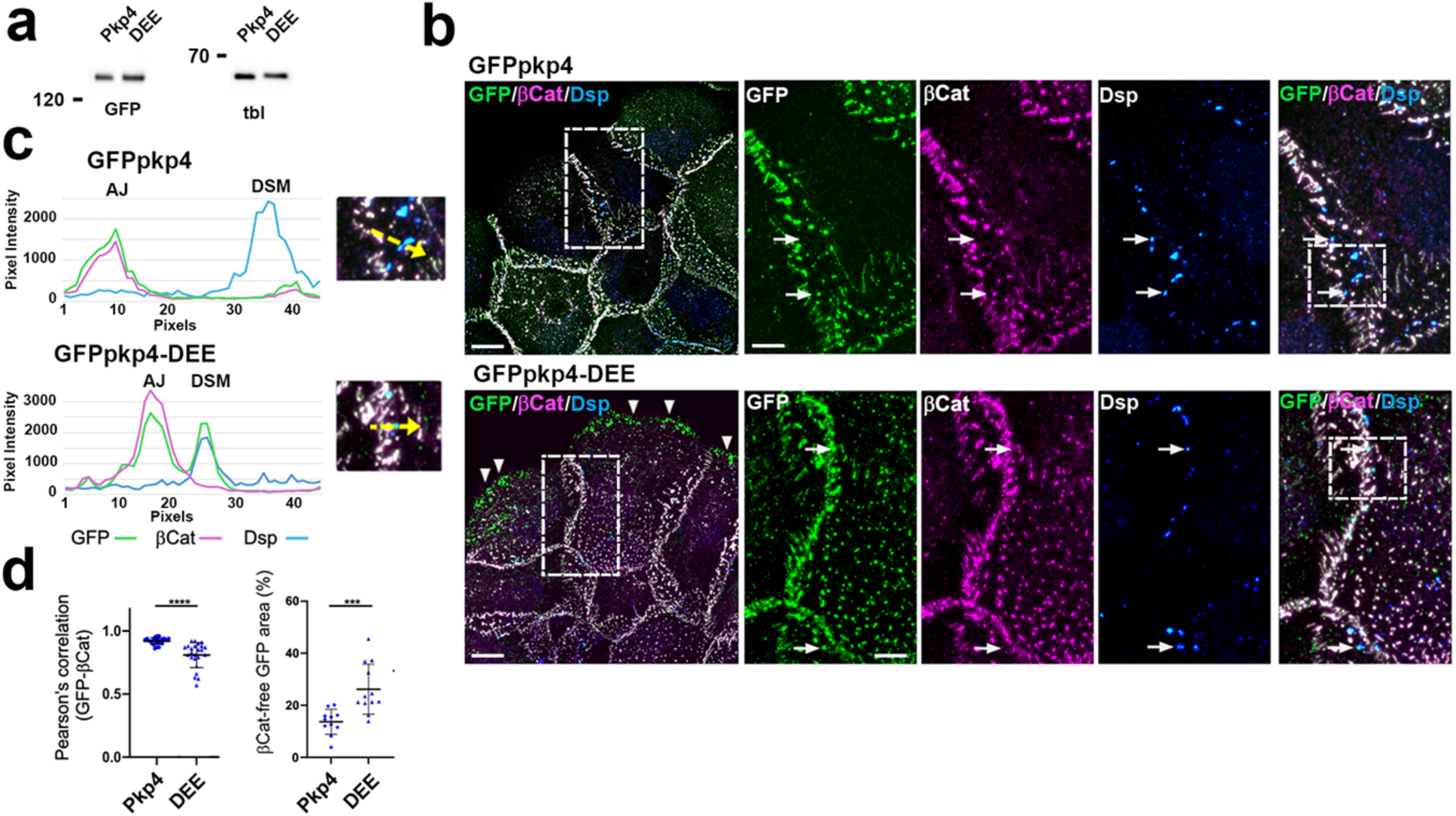
Phospholipid-uncoupled Pkp4 mutant localizes to AJs through the cadherin-dependent pathway. (**a**) Immunoblot analysis of A431-δCat-KO cells expressing GFPpkp4 (Pkp4) or its phospholipid-uncoupled mutant, GFPpkp4^DEE^ (DEE). Blots were probed for GFP and β-tubulin (tbl; loading control). Molecular mass markers (kDa) are shown on the left. (**b**) Maximum-intensity projections of confocal z-stacks from cells expressing GFPpkp4 or GFPpkp4^DEE^ (GFPpkp4-DEE). Cells were stained for GFP (green), β-catenin (βCat, magenta), and desmoplakin (Dsp, blue). Representative cell-cell contact regions (dashed boxes) are enlarged on the right and shown as individual fluorescence channels. Scale bars, 10 μm (main images) and 4 μm (enlargements). Disruption of phospholipid binding by the DEE mutation does not noticeably affect localization of Pkp4 to AJs. However, two abnormalities are evident. First, desmosomes (arrows), which do not recruit wild-type GFPpkp4, efficiently recruit GFPpkp4^DEE^. Second, the mutant accumulates in large intracellular aggregates (arrowheads), particularly along the free edge of the cell islands. (**c**) Fluorescence intensity profiles measured along the dashed yellow lines shown in the magnified regions outlined by the dashed boxes in (**b**). The profiles illustrate selective recruitment of wild-type GFPpkp4 to AJs, whereas GFPpkp4^DEE^ localizes to both AJs and desmosomes. (**d**) Pearson’s correlation coefficients between GFP and β-catenin fluorescence (left) and the abundance of extrajunctional GFP signal (right), quantified as the fraction of GFP-positive pixels lacking β-catenin staining relative to the total GFP-positive pixels. Consistent with the formation of intracellular aggregates and desmosomal localization, this parameter is significantly higher in GFPpkp4^DEE^ relative to GFPpkp4 cells. Statistical significance was calculated using two-tailed Student’s t tests: ***<P 0.001; ****P < 0.0001. The means ± SD are indicated by bars.

### Direct interactions with E-cadherin are dispensable for recruitment of endogenous Pkp4, but not p120, to AJs

Our experiments demonstrated that Pkp4 can be targeted into AJs independently of its direct binding to classical cadherins. However, these studies relied exclusively on overexpressed GFP-tagged Pkp4. Under physiological conditions, Pkp4 is expressed at much lower levels than its closely related family member p120 and therefore competes with p120 for incorporation into AJs. To determine whether phospholipid-dependent recruitment of Pkp4 into AJs also operates under physiological Pkp4 expression level, we examined the endogenous Pkp4 upon its uncoupling from cadherins. To abolish direct δ-catenin-cadherin interactions, we expressed in cadherin-deficient A431-Ec/Pc-KO cells a GFP-tagged E-cadherin mutant, EcGFP-KLΔJMD, lacking the core δ-catenin-binding site (JMD) together with its adjacent flanking region (Fig. 7a). To stabilize this mutant at the plasma membrane, its two endocytic motifs were simultaneously inactivated by the K738R and L741V/L742A substitutions (Hong et al., 2010). The resulting mutant was expressed at levels comparable to those of endogenous E-cadherin in wild-type A431 cells and intact EcGFP in A431-Ec/Pc-KO cells (Fig. 7b). Immunoblotting further confirmed that EcGFP- and EcGFP-KLΔJMD-expressing cells contained similar levels of their endogenous p120 and Pkp4 (Fig. 7b).

**Figure 7.**
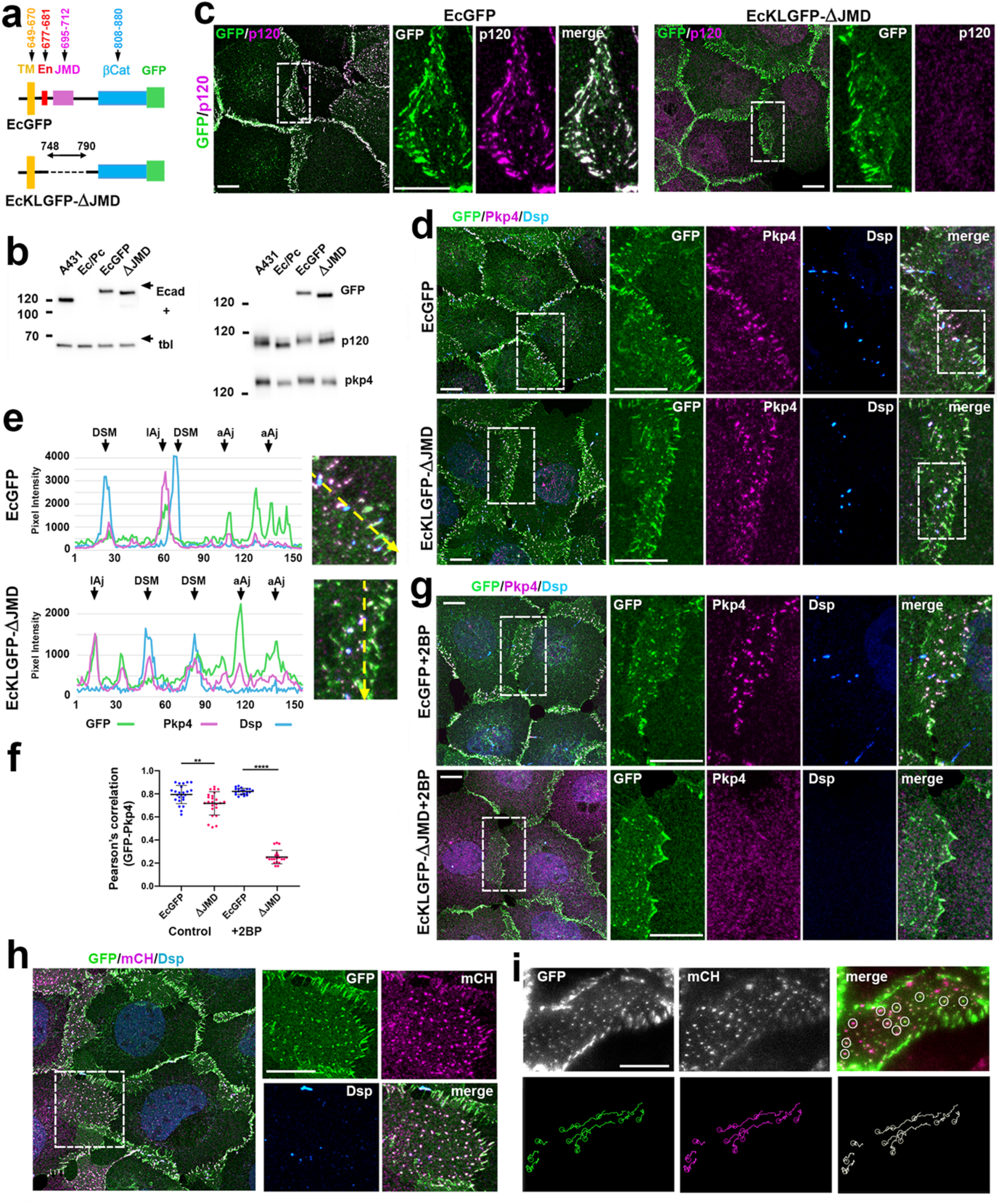
δ-Catenin-binding site of E-cadherin is required for p120 but not for Pkp4 recruitment into AJs. (**a**) Schematic representation of the C-terminal region of EcGFP and its δ-catenin-uncoupled mutant, EcKLGFP-ΔJMD. The EcGFP contains the transmembrane domain (TM, orange), the juxtamembrane δ-catenin-binding domain (JMD, magenta), and the β-catenin-binding region (βCat, blue) as well as two endocytic motifs: the ubiquitination site, K738, and the dileucine motif, L741/L742 (En, red). EcKLGFP-ΔJMD lacks both endocytic motifs and also the entire region upstream of the β-catenin-binding site that encompasses the JMD. (**b**) Immunoblot analysis of wild type A431 cells (A431), their cadherin deficient variant, A431-Ec/Pc-KO subline (Ec/Pc), and its derivatives expressing EcGFP and EcKLGFP-ΔJMD (ΔJMD). Blots were probed for E-cadherin (Ecad), β-tubulin (tbl), GFP, p120, and Pkp4. Note that the GFP-tagged proteins are expressed at levels comparable to endogenous E-cadherin. Molecular mass markers (kDa) are shown on the left. (**c**) Maximum-intensity projections of confocal z-stacks from cells expressing EcGFP and EcKLGFP-ΔJMD stained for GFP (green) and p120 (magenta). Representative cell-cell contact regions (dashed boxes) are enlarged on the right and shown as individual fluorescence channels. Scale bars, 10 μm. Note that JMD deletion in E-cadherin completely abolished p120 association with AJs. (**d**) Maximum-intensity projections of confocal z-stacks from the same cells as in (c), stained for GFP (green), Pkp4 (magenta), and Dsp (blue), and displayed as in (c). In contrast to p120, Pkp4 remains associated with lateral AJs and additionally localizes to desmosomes in EcKLGFP-ΔJMD cells. (**e**) Fluorescence intensity profiles measured along the dashed yellow lines in the regions outlined by the dashed boxes in (d). The profiles illustrate selective localization of endogenous Pkp4 to lateral AJs in EcGFP cells and to both lateral AJs and desmosomes in EcKLGFP-ΔJMD cells. (**f**) Pearson’s correlation coefficients for GFP and Pkp4 fluorescence within cell-cell contacts of cells expressing EcGFP or EcKLGFP-ΔJMD under control conditions and after treatment with 2-bromopalmitate (+2BP). Statistical significance was calculated using two-tailed Student’s t tests: **<P 0.01; ****P < 0.0001. The means ± SD are indicated by bars. (**g**) Projections of confocal z-stacks from the same cells as in (c) following 2BP treatment. Cells were stained and displayed as in (**c**). (**h**) EcKLGFP-ΔJMD cells expressing mCherry-tagged Pkp4 (mCHpkp4) were stained for GFP (green) and mCherry (magenta) and displayed as in (**c**). Note, that recombinant Pkp4 preferentially incorporates into lateral AJs despite the absence of the cadherin JMD. (**i**) Initial frames from the green (GFP) and red (mCherry) channels of a ∼1h time-lapse recording (Video 1) of EcKLGFP-ΔJMD cells co-expressing mCHpkp4. Representative lateral AJs (circles) that could be tracked throughout the recording are shown below (circles display their initial positions). Scale bar, 10 μm. Note, that the GFP- and mCherry-tagged proteins remain tightly associated throughout the entire recording.

Since endogenous classical cadherins are absent from A431-Ec/Pc cells, all AJs in EcGFP-KLΔJMD-expressing cells are formed by the mutant E-cadherin and therefore lack the canonical δ-catenin-binding site. As expected, p120 was readily detected in nearly all AJs of EcGFP-expressing cells but was undetectable in the AJs of EcGFP-KLΔJMD cells (Fig. 7c). In striking contrast, and in complete agreement with our experiments using recombinant Pkp4, endogenous Pkp4 remained associated with the majority of lateral AJs, exhibiting only a modest reduction in its PCC with E-cadherin (Fig. 7d). Furthermore, whereas Pkp4 in EcGFP-expressing cells localized almost exclusively to AJs, EcGFP-KLΔJMD cells contained numerous additional Pkp4-positive clusters at cell-cell contacts. Co-staining with desmoplakin identified these structures as desmosomes (Fig. 7d,e), while staining for the lateral AJ marker, PLEKHA5, confirmed that the remaining Pkp4-positive adherens junctions represented lateral AJs (Fig. 7g). Thus, the behavior of endogenous Pkp4 fully recapitulated that observed with the GFP-tagged Pkp4^WN^ mutant, demonstrating that even under physiological expression level, the direct binding of Pkp4 to the cadherin JMD is not required for its recruitment to lateral AJs. At the same time, loss of the canonical δ-catenin-binding site resulted in increased accumulation of Pkp4 in desmosomes, suggesting that direct cadherin binding normally contributes to restricting Pkp4 to AJs.

Our previous studies showed that individual lateral AJs can be monitored in living cells as highly stable, mobile dot-like structures over periods of at least one hour (Indra et al., 2026). To determine whether this exceptional stability depends on direct Pkp4-cadherin interactions, we analyzed dynamics of the EcGFP-KLΔJMD-containing AJs in A431-Ec/Pc-KO cells co-expressing mCherry-tagged Pkp4^WNHR^ mutant (Fig. 7h, see also Fig. S1). Remarkably, despite the absence of direct Pkp4-cadherin binding, both mCherry-Pkp4 ^WNHR^ and EcGFP remained stably associated within the lateral AJs throughout the entire 1-hour imaging period, exhibiting the characteristic oscillatory movements of these junctions (Fig. 7i; Movie S1). Together, these experiments confirm that, unlike p120, the endogenous Pkp4 can generate exceptionally stable lateral AJs through a mechanism that is independent of the direct binding to the cadherin JMD.

## Discussion

δ-Catenins, members of the ARM-repeat protein superfamily, interact with both classical and desmosomal cadherins and play still poorly understood roles in the assembly and regulation of cadherin-mediated cell-cell adhesion. The major classical cadherin-associated δ-catenin, p120, is dispensable for the core adhesive function of AJs, which primarily depends on α-catenin. The dispensability of p120 in AJs is supported by genetic studies in invertebrates (McCrea & Park JI, 2007), by experiments using cadherin-α-catenin chimeras (Hong et al., 2013, Troyanovsky et al., 2015), and by the observation that adhesion defects caused by p120 depletion in vertebrate cells can largely be rescued by preventing cadherin endocytosis (Ireton et al., 2002; Schell et al., 2026). On the other hand, accumulating evidence indicates that δ-catenins are indispensable players in the assembly and function of specialized cadherin-based junctions, including desmosomes, neuronal synapses, and cardiac intercalated discs. Their importance is underscored by the numerous human skin, neurological, and cardiac disorders caused by mutations in δ-catenin family members (Kosik et al., 2005; McCrea & Park, 2007; Bass-Zubek et al., 2009; Hofmann, 2020; Donta et al., 2022). Functional studies further support this conclusion. For example, among the major structural components of desmosomes, only plakophilins are absolutely required for desmosome assembly (Todorovic et al., 2014; Fujiwara et al., 2015; Indra et al., 2021). Despite compelling genetic and functional evidence of their importance, the mechanisms by which plakophilins support the assembly of these junctions remain largely unknown. Here, we report two unexpected findings that provide new mechanistic insights into how δ-catenins regulate cadherin-based adhesions.

We focused our study on Pkp4 because our recent work identified this δ-catenin as the key organizer of a specialized subtype of AJs, termed lateral AJs. Although these junctions incorporate classical cadherins and are widely distributed in epithelial tissues (Takeichi, 2014; Franke et al., 2009; Hofmann et al., 2008), we have shown that they are assembled through α-catenin-independent mechanism and appear to play a signaling rather than a primary adhesive role (Indra et al., 2026). Here, we challenge the prevailing view that AJ-associated δ-catenins are recruited exclusively through direct binding to the cadherin JMD (Anastasiadis & Reynolds, 2000): we show that Pkp4 is recruited to lateral AJs independently of its canonical interaction with the cadherin JMD. This conclusion is supported by two alternative strategies: disruption of the JMD-binding interface in Pkp4 and deletion of the JMD from E-cadherin. In both cases, Pkp4 remained tightly associated with lateral, but not apical or basal, AJs despite the absence of direct cadherin binding. Moreover, even in cells completely lacking classical cadherins and, therefore, unable to assemble AJs, Pkp4 still formed trans-interacting clusters at sites of cell-cell contacts. Together, these findings suggest that Pkp4 can be targeted to lateral AJs through a positional cue generated at sites of cell-cell contacts independently of classical cadherins. The ability of a cadherin-uncoupled Pkp4 mutant lacking both the N- and C-terminal domains to localize to lateral AJs further suggests that this targeting is mediated by the conserved ARM domain of Pkp4. The N- and C-domains are apparently implicated in Pkp4 oligomerization at the sites of targeting, the process that requires a separate study.

We also identified the molecular basis of the positional cue. It depends on interactions between ARM domain of Pkp4 and phospholipid-rich plasma membrane domains. Specifically, we found that the polybasic motif located within the insert separating the ARM repeats 5 and 6 (5/6INT^pkp4^) binds several plasma membrane phospholipids, including PI(4,5)P2, which is known to be involved in AJ formation (Ling et al., 2007; Xiong et al., 2012) and in specialization of the lateral membrane of epithelial cells (Kanematu et al., 2022). Interestingly, the proximity proteomics presented in the latter work showed the Pkp4, but not p120, is one of the PI(4,5)P2-proximal proteins. Furthermore, the disruption of phospholipid binding abolished cadherin-independent targeting of Pkp4 to lateral AJs, demonstrating that this interaction is essential for this pathway. The polybasic motif within the 5/6INT^pkp4^ is adjacent to the δ-catenin conserved cysteine residue (Cys785 in Pkp4), which has previously been shown to be palmitoylated in desmosomal plakophilins (Roberts et al., 2014). Mutation of this conserved cysteine produced essentially the same phenotype as disruption of the polybasic motif, abolishing cadherin-independent targeting of Pkp4 to cell-cell contacts. Consistent with this conclusion, pharmacological inhibition of palmitoylation with 2-bromopalmitate also abolished cadherin-independent targeting of Pkp4.

Taken together, results reported here show that Pkp4 could be targeted to cell-cell junctions by two complementary and independent mechanisms. One is mediated by the canonical interaction with classical cadherins, whereas the other depends on recognition of phospholipid-rich plasma membrane domains. The relative contribution of these two mechanisms determines where Pkp4 is incorporated. When Pkp4 is overexpressed and efficiently competes with p120 for cadherin binding, it is incorporated broadly into all available AJs. However, under physiological conditions, where Pkp4 is a minor δ-catenin, it preferentially targets CCCs to lateral AJs since this targeting is reinforced by cooperation of two mechanisms.

The conservation of the 5/6INT throughout the δ-catenin family, including the conserved palmitoylated cysteine, strongly suggests that recognition of phospholipid-rich membrane is a common property of δ-catenin ARM domains. This conclusion is consistent with the established role of lipid rafts and the 5/6INT palmytoilation in desmosome biogenesis (Roberts et al., 2014; Zimmer et al., 2024; Zimmer & Kowalczyk, 2024), as well as with the observation that alternative splicing of p120, which disrupts the polybasic 5/6INT motif, causes cytosolic accumulation of the corresponding isoform (Troyanovsky et al 2011).

The interactions between δ-catenins and phospholipid-rich domains are likely to be utilized in different ways by distinct δ-catenins and in different cellular contexts. For example, in AJs, this interaction may contribute to the local regulation of cadherin endocytosis. The major cargo endocytic adaptor, AP-2 (Chiasson et al., 2009), recognizes cargo endocytic motifs only upon binding to phospholipids (Sloan et al., 2026). An attractive possibility, therefore, is that δ-catenins compete with AP-2 for membrane phospholipids, thereby suppressing cadherin endocytosis. This model is consistent with the selective destabilization of lateral and apical/basal AJs following the loss of Pkp4 and p120, respectively (Indra et al., 2026). Association of δ-catenins with phospholipid-rich domains may also couple AJs to several key membrane-associated regulatory systems, including the apicobasal polarity machinery and regulators of the actin cytoskeleton, both of which are strongly influenced by local membrane lipid composition (Teo et al., 2019; Thapa et al., 2023; Senju & Lappalainen, 2019). In desmosomes, this property may play an even more fundamental role by defining the membrane sites that permit clustering of desmosomal cadherins.

In conclusion, our data show that the ARM domain of δ-catenins integrates two independent binding activities: direct recognition of cadherins and recognition of phospholipid-rich membrane domains. The differences between δ-catenin family members in the specificity and relative contribution of these two activities could provide a general mechanism for generating the remarkable structural and functional diversity of cadherin-based junctions across vertebrate tissues.

## Materials and Methods

### Plasmids

The original plasmid, pRcCMV-GFPpkp4, containing a GFP-tagged full-length Pkp4 cDNA, was previously described (Indra et al., 2026). The mutants of this plasmid, as well as its mCherry version were constructed using PCR-based mutagenesis. The general maps of the mutant are presented in Figs. 1a and 5a. Before the use, all plasmids were verified by the full plasmid sequencing. The DNA fragment encoding the GFP-CC-5/6INT^PKP4^ biosensor (synthesized by Integrated DNA Technologies) was subcloned downstream to a GFP-encoding portion of the pRcCMVGFP-αCat plasmid (Chen et al., 2015). The resulting biosensor contained GFP-tagged antiparallel coiled-coil domain of seryl-tRNA synthetase from Thermus thermophilus (Amer et al., 2023), which loop, connecting two helices, was replaced with the central portion of the 5/6 insert of Pkp4 (residues 761-811). The resulting amino acid sequence of the biosensor is: ggssvfvvaerellaldrevqelkkrlqevqternqvakrvgRLLGLNELDDLLGKESPSKDSEPSCWGKKKK KKKRTPQEDQWDGVGPIPGLgliargkalgeeakrleealrekearlealrgehralgsgstg, where the uppercase letters represent the 5/6 insert and lowercase – the coiled-coil helices.

### AlphaFold 3 structural modeling

Structural modeling of the complexes of Pkp4-ARM or its mutants with E-cadherin JMD was performed using the AlphaFold 3 (alphafoldserver.com). The models were generated for Pkp4-ARM (residues 524-1026) or corresponding Pkp4-ARM-WNHR mutant in complex with E-cadherin JMD (residues 695-712), and for the full-length GFP-CC-5/6INT^PKP4^ sequence. The resulting models were visualized by the Web-based 3D Structure Viewer, Icn3D.

### Cell culture and transfection

The original A431 cell line and its derivates deficient in classical cadherins (A431-Ec/Pc-KO), in Pkp4 (A431-Pkp4-KO), and in Pkp4/p120/ARVCF (δCat-KO-A431) have been previously described (Troyanovsky et al., 2021; Indra et al., 2026). The cells were grown in DMEM supplemented with 10% FBS and were transfected using Lipofectamine 2000 (Invitrogen) according to the company protocol. After selection of the Geneticin-resistant cells (0.5 mg/ml), the cells were sorted for transgene expression by FACS, and only moderate-expressing cells were used. At least three clones were selected for each construct, and all were tested in most of the assays. The expression levels and sizes of the recombinant proteins in the obtained clones were analyzed by Western blotting as previously described (Troyanovsky et al., 2025) using anti-GFP or anti-mCherry antibodies and Nitrocellulose Blotting Membrane (Amersham, #10600007). All clones of cells expressing a particular transgene exhibited the same phenotype. Representative data for one of three clones is presented. The drugs, GSK-A1 (MedChemExpress, HY-125118) and 2-bromopalmitic acid (Sigma, 238422) were used at concentration 100 nM for 30 min and 50 μM for 5 h, correspondingly.

### Phospholipid binding and actin co-sedimentation

For recombinant GST fusion protein production, the Pkp4 DNA fragment encoding its ARM domain (residues 509-1025) was subcloned into the bacterial expression vector pGEX-4T-1, which places GST in front of the Pkp4 fragment. The resulting plasmids were transformed into BL21(DE3) cells. Protein isolation was performed essentially as described (Laur et al 2002; Chen et al., 2015). In brief, the cells (induced for protein expression using 1 mM isopropyl β-D-1-thiogalactopyranoside) were harvested by centrifugation (5000×g, 25 min) and lysed by sonication in lysis buffer (50 mM Tris, pH 8, 150 mM NaCl). The lysate was clarified (35,000 rpm, 30 min) and the proteins were purified using GST Spin Purification kit (Pierce) with no deviation from the manufacturer’s protocol. The purity of the products was tested in SDS-PAGE, and protein concentration was estimated using “reducing agent compatible BSA protein assay kit” (Thermo Fisher Scientific, 23250). The co-sedimentation assay was performed as described previously (Chen at al., 2015) using GST-αABD as a positive control. In brief, the pre-polymerized actin filaments (rabbit skeletal muscle, Cytoskeleton, Inc.) in F buffer (2 mM Tris, pH 8.0, 50 mM KCl, 2 mM MgCl2, 1 mM ATP, 1 mM EGTA, and 1 mM DTT) were incubated with precleared (100,000 g for 30 min) recombinant proteins for 30 min at room temperature. The samples were then centrifuged at 100,000 g for 30 min. Equivalent volumes of pellet and supernatant fractions were analyzed by SDS-PAGE, stained with Coomassie Brilliant Blue, scanned on Azure C300 Chemiluminescent Imager (Azure Biosystems, Dublin, CA), and quantified with FIJI. The protein-lipid overlay assay was performed using the Mega Lipid Strips (Echelon Bioscinces, P-6005) according to the company protocol. In brief, the strips were subsequently incubated (for 1h each time) with: (i) blocking buffer, (ii) GST-tagged proteins, at concentration 0.5 nM, (iii) mouse anti-GST antibody (1:2000 of original concentration, Invitrogen, 3215837), (iv) goat peroxidase-conjugated anti-mouse antibody (1:1000, Cell Signaling, 7076). Chemiluminescence was detected via the Azure C300.

### Immunofluorescence microscopy

For immunofluorescence, cells were grown for 2 days on glass coverslips and were fixed with 3% formaldehyde (5 min) and then permeabilized with 1% Triton X-100 (15 min), as described previously (Indra et al., 2020; Klingelhöfer et al., 2000). The confocal images were taken using the Nikon AXR laser scanning microscope equipped with a Plan Apo 60x×/1.45 objective lens. The images were processed using Nikon’s NIS-Elements software. For immunostaining and Western blotting the following antibodies were used: Mouse anti-E-cadherin mAb clones SHE78-7 and HECD1 (Takara, M126 and M106), anti-GFP (Santa Cruz Biotechnology, sc-9996), anti-Pkp4 (Progen, 651166), anti-GFP (Invitrogen, A11120) and anti-p120 (BD Transduction Laboratories, 610134); Chicken anti-GFP (Novus, NB100-1614); Rabbit anti-p120 (Abcam, ab92514), anti-Dsg2 (Proteintech, 21880-I-AP), anti-mCherry (BioVision, 5993-100), anti-PLEKHA5 and anti-Pkp4 (Invitrogen, PA5-57463 and PA5-66855, respectively); Guinea pig anti-Pkp4 and anti-desmoplakin (Progen, GP71 and DP-1). Donkey Alexa Fluor 488-conjugated anti-rabbit (711-545-152), anti-mouse (715-545-150), anti-goat (705-545-147), anti-guinea pig (706-545-148); donkey Cy3-conjugated anti-rabbit (711-165-152), anti-mouse (715-165-150), anti-goat (705-165-147), anti-guinea pig (706-165-148); Alexa Fluor 647-conjugatred donkey anti-guinea pig (706-605-148); and donkey horseradish peroxidase-conjugated anti-mouse (715-035-150), anti-rabbit (711-035-152), anti-guinea pig (706-035-148) secondary antibodies were purchased from Jackson Immunoresearch Laboratories.

### Live-cell imaging

The live cell imaging experiments were performed essentially as described previously (Indra et al., 2018) using an X-Cite 120LED Boost High-Power LED Illumination System as a light source. In brief, cells were imaged in L-15 media with 10% FBS by an Eclipse Ti-E microscope (Nikon, Melville, NY) at RT or 37°C controlled with Nikon’s NIS-Elements software. No binning mode was used. At this microscope setting, the pixel size was 110 nm. To monitor the entire lateral membrane from its basal to the apical edges, the stacks of 5 focal planes with 0.5 μm spacing were taken at each time point. All z-stack images were saved in ND2 format. The maximum intensity projection of each frame was created using NIS Element 5.02. For movie analyses, all images were saved as Tiff files and processed using ImageJ software (National Institutes of Health). Semi-automatic tracking of lateral AJs was performed using the TrackMate toolbox in Fiji. For each dataset, ten randomly selected lateral AJs were chosen for tracking. Individual puncta were initialized by selecting the region of interest, and the selection size was adjusted as needed ensuring that the punctum was accurately encompassed. The selected puncta were then tracked over time using the semi-automatic tracking workflow in TrackMate. All generated tracks were visually inspected for accuracy throughout the time series.

### Electron Microscopy

Cells grown on glass coverslips were sequentially extracted for 5 min with cytoskeleton preservation buffer (CPB, 100 mM PIPES, pH 6.9; 1 mM EGTA; 1 mM MgCl2) supplemented with 1% Triton X-100, then washed with CPB for 1 min, and incubated for 10 min with the mouse anti-GFP antibody (Invitrogen) in CBP. Then the cells were washed for 5 min in CBP and incubated for additional 10 min with Biotin-SP-VHH fragment Alpaca anti-mouse antibody (615-064-214, Jackson ImmunoResearch Laboratories). The staining solutions were supplemented with 1% of Blocking Solution for Gold conjugates (Aurion). After another washing, the cells were fixed with 0.2% glutaraldehyde (for 10 min), quenched with 2 mg/ml NaBH_4_ in PBS (for 10 min) and stained with 10 nm gold-conjugated Streptavidin (Abcam, ab270041). After final washing cells were fixed with 2% glutaraldehyde in 0.1 M Na-cacodylate, pH 7.3. After post-fixation in 1% osmium tetroxide and 3% uranyl acetate, samples were dehydrated in series of ethanol, embedded in Epon812 resin, and polymerized for 48 hours at 60°C. Then ultrathin sections were made using Ultracut UC7 Ultramicrotome (Leica Microsystems, Germany), which were contrasted with 3% uranyl acetate and Reynolds lead citrate. A parallel coverslip (stained as described above) were fixed in 3% formaldehyde and stained with rabbit anti-Streptavidin antibody to verify staining specificity. EM samples were imaged using a FEI Tecnai Spirit G2 transmission electron microscope (FEI Company, Hillsboro, OR) operated at 80 kV. Images were captured by Eagle 4k HR 200kV CCD camera and presented in inverted contrast.

### Data processing

Chemiluminescence and Coomassie staining was recorded via the Azure C300 Chemiluminescent Imager (Azure Biosystems, Dublin, CA) and band intensities were analyzed using ImageJ software (rsb.info.nih.gov/ij/). Most cell images were processed and analyzed using Nikon’s NIS-Elements ver. 5.02. For line scan analysis, the Element’s in-built line profile function was used to draw a 1pix wide line across the representative cell-cell contact areas. To assess the ratio of the monochromatic GFP-positive pixels versus the total number of the GFP-positive pixels (Fig. 6d), channels were split and treated with median filter of 1 pixel using Fiji. Both channels were auto-thresholded using Otsu algorithm. Thresholds were selected and added to the ROI manager. ‘AND’ function was used to include the overlapping area from both channels. Percentage of overlap was calculated from the resulting analysis. To quantify the GFP and β-catenin signal at apical and lateral punctate junctions, fluorescence intensities of GFP and β-catenin were measured within a standardized region of interest (ROI) of 400 µm². Separate ROIs were selected for apical and lateral punctate junctions, and the fluorescence intensity of each channel was quantified within the same ROI. The GFP/βCat index (Fig. 1g) was calculated by dividing the measured GFP fluorescence intensity by the corresponding β-catenin fluorescence intensity for each 400 µm² ROI. For Pearson’s correlation quantification, the arbitrary selected representative cell-cell contact areas were processed with limited background reduction and denoising function of NIS-Element 5.02 and then Element’s colocalization tool was used to obtain the scatterplots of red and green pixel intensities and to evaluate the Pearson’s correlation coefficient (PCC). PCC of representative 5 cell-cell contact areas taken from three independent images were quantified (Figs 1e, 3e, 4c, 6d, 7f). The charts and error bars were plotted using GraphPad Prism version 10.2.0. Statistical significance was analyzed using student’s two-tailed t test for two groups. A p value that was less than 0.05 was considered statistically significant.

## Acknowledgments

We thank Dr. R. Leube (RWTH Aachen University) for valuable comments and suggestions. Sequencing, flow cytometry and confocal microscopy were performed at the Northwestern University Genetic, Flow Cytometry, and Advanced Microscopy Centers. The authors declare no competing financial interests. The work was supported by National Institute of Health Grant AR070166 (to S.M.T.).

---

**Figure S1.**
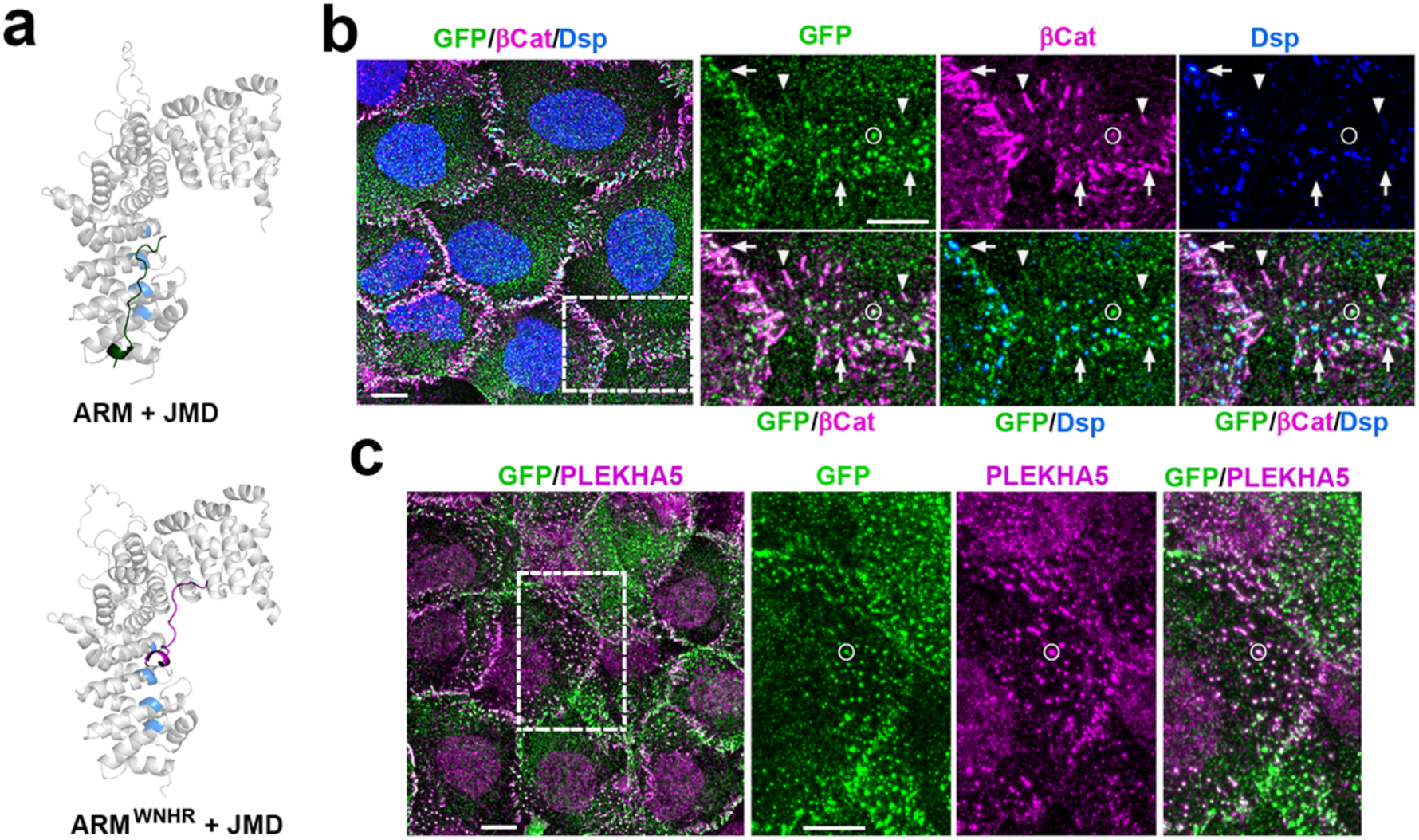
Cadherin-uncoupled Pkp4 preferentially associates with lateral AJs. (**a**) AlphaFold 3-generated structural models of the E-cadherin JMD bound to the Pkp4 ARM domain or its WNHR mutant. The model generated for the wild-type ARM domain (ARM+JMD) shows that the positively charged groove (blue) of the Pkp4 ARM domain (gray) accommodates the JMD (green). In the WNHR mutant (ARMWNHR+JMD), this groove no longer recognizes the JMD, which instead binds to an alternative, nonphysiological interface (red). (**b**, **c**) Maximum-intensity projections of confocal z-stacks from cells expressing GFPpkp4^WNHR^ and stained (**b**) for GFP, β-catenin (βCat), and desmoplakin (Dsp), or (**c**) for GFP and PLEKHA5. Boxed regions are enlarged on the right. Scale bar, 10 μm. Note, that GFPpkp4WNHR is preferentially recruited to lateral AJs, identified by β-catenin but lacking Dsp in (**b**) and by PLEKHA5 in (**c**), as well as to desmosomes, identified by Dsp but lacking β-catenin. Representative lateral AJs are indicated by circles. In contrast, apical (arrows) and basal (arrowheads) AJs exhibit markedly reduced GFP labeling.

**Figure S2.**
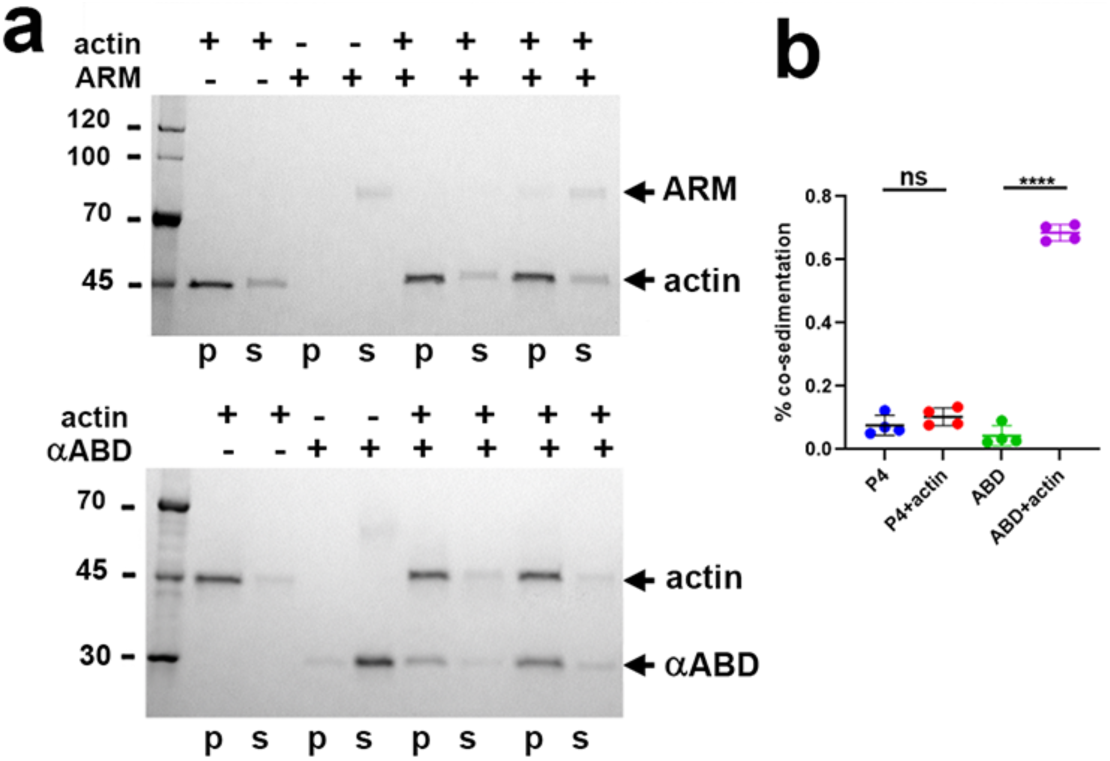
Pkp4 ARM domain exhibits no detectable interactions with F-actin. (**a**) Representative F-actin co-sedimentation assay. SDS-PAGE analysis of the pellet (p) and supernatant (s) fractions obtained after incubation of F-actin with GST-tagged Pkp4 ARM domain (ARM; top gel) or GST-tagged α-catenin actin-binding domain (αABD; positive control, bottom gel). GST-αABD efficiently co-sediments with F-actin, whereas GST-ARM^PKP4^ remains predominantly in the supernatant. (**b**) Quantification of the co-sedimentation assay showing that, in contrast to GST-αABD, GST-ARM^PKP4^ exhibits no detectable binding to F-actin. Data are from n = 4 independent experiments. Statistical significance was calculated using two-tailed Student’s t tests: ns, nonsignificant; ****P < 0.0001. The means ± SD are indicated by bars.

**Figure S3.**
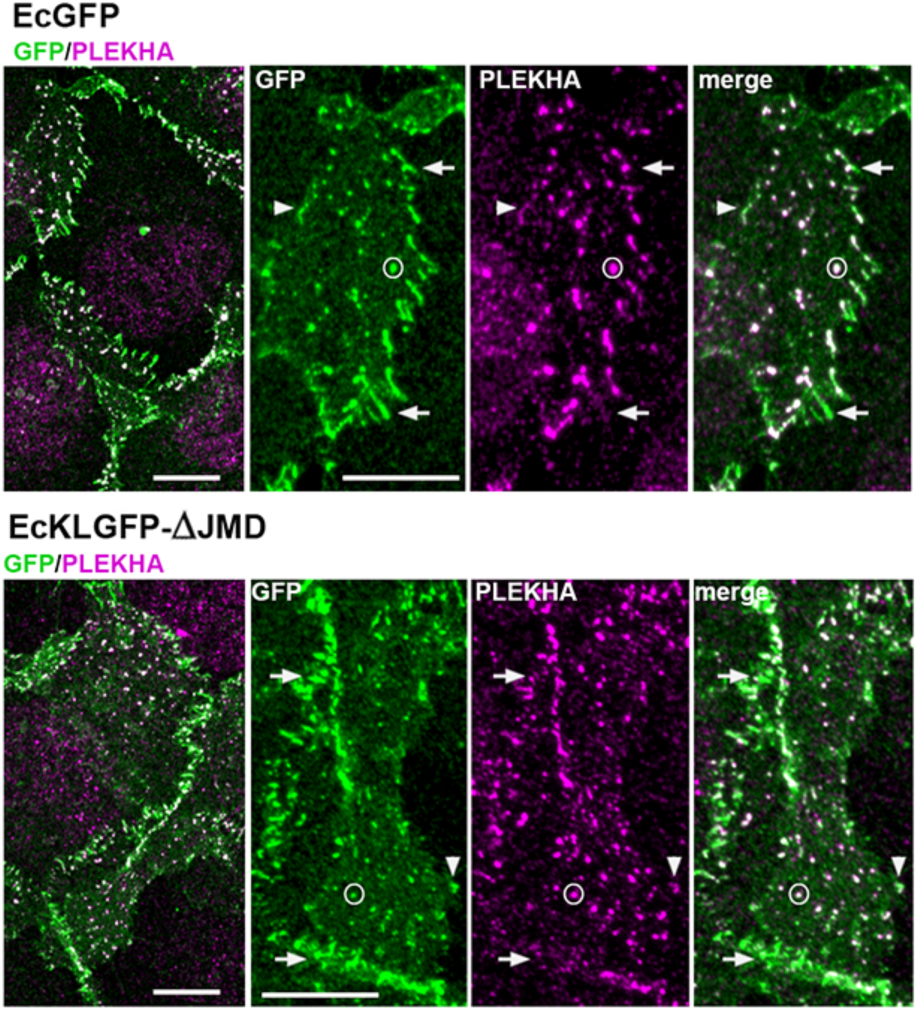
δ-Catenin-uncoupled E-cadherin mutant forms lateral AJs. A431-Ec/Pc-KO cells expressing EcGFP (positive control) or the δ-catenin-uncoupled E-cadherin mutant EcKLGFP-ΔJMD were stained for GFP (green) and the lateral AJ marker PLEKHA5 (magenta) and analyzed by confocal microscopy. Representative cell-cell contact regions (dashed boxes) are enlarged on the right and shown as individual fluorescence channels. Scale bar, 10 μm. Note, that both cell lines recruit PLEKHA5 preferentially to lateral AJs (representative junctions are indicated by circles), but not to apical or basal AJs (examples are indicated by arrows and arrowheads, respectively).

**Video S1. Joint dynamics of Pkp4 and Pkp4-uncoupled E-cadherin mutant, EcKLGFP-ΔJMD, at the selected cell-cell contact.** Time-lapse of AJs in Ec/Pc-KO cells co-expressing EcKLGFP-ΔJMD (green) and mCHpkp4 (red). The most representative cell-cell contact from 6 independent movies was selected. The images were acquired at 30 sec intervals and are shown with display rate 10 frames/sec. See Fig. 7 for details.

